# High-content analysis of proteostasis capacity in cellular models of amyotrophic lateral sclerosis (ALS)

**DOI:** 10.1101/2021.09.23.461468

**Authors:** Isabella A. Lambert-Smith, Justin J. Yerbury, Darren N. Saunders

## Abstract

Disrupted proteome homeostasis (proteostasis) in amyotrophic lateral sclerosis (ALS) has been a major focus of research in the past two decades. Yet the exact processes that normally maintain proteostasis, but that are uniquely disturbed in motor neurons expressing diverse genetic mutations, remain to be established. Obtaining a better understanding of proteostasis disruption in association with different ALS-causing mutations will improve our understanding of ALS pathophysiology and may identify novel therapeutic targets and strategies for ALS patients. Here we describe the development and use of a novel high-content analysis (HCA) assay to investigate proteostasis disturbances caused by the expression of ALS-causing gene variants. This assay involves the use of conformationally-destabilised mutants of firefly luciferase (Fluc) to examine protein folding/re-folding capacity in NSC-34 cells expressing ALS-associated mutations in the genes encoding superoxide dismutase-1 (SOD1^A4V^) and cyclin F (CCNF^S621G^). We demonstrate that these Fluc isoforms can be used in high-throughput format to report on reductions in the activity of the chaperone network that result from the expression of SOD1^A4V^, providing multiplexed information at single-cell resolution. In addition to SOD1^A4V^ and CCNF^S621G^, NSC-34 models of ALS-associated TDP-43, FUS, UBQLN2, OPTN, VCP and VAPB mutants were generated that could be screened using this assay in future work. For ALS-associated mutant proteins that do cause reductions in protein quality control capacity, such as SOD1^A4V^, this assay has potential to be applied in drug screening studies to identify candidate compounds that can ameliorate this deficiency.

**Graphical abstract:** 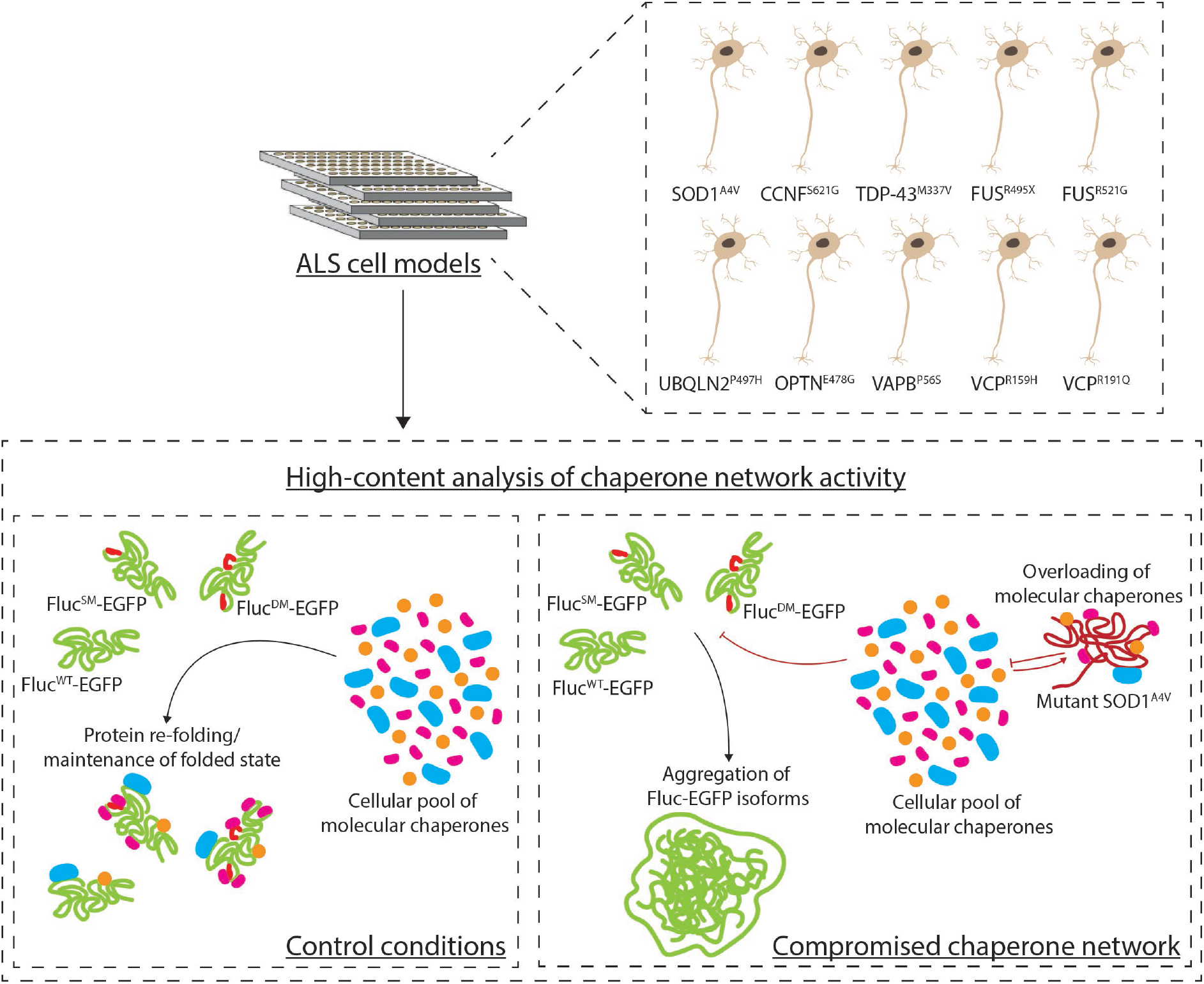

**Highlights:**

- Destabilised firefly luciferase (Fluc) mutants can be used in high-content analysis (HCA) assay format
- Fluc HCA assay enables information-rich reporting on chaperone network activity in cell models of ALS
- Expression of SOD1^A4V^ reduces chaperone network activity in NSC-34 cells

## 1. INTRODUCTION

Post-mortem examination of spinal cord tissue from amyotrophic lateral sclerosis (ALS) patients consistently reveals the presence of TDP-43-, FUS- or SOD1- and ubiquitin-positive inclusions comprised of insoluble proteinaceous material [1-4]. Misfolded proteins, as either abnormal monomers and/or oligomeric precursors, possess cytotoxic properties [5-7] and their aggregation into certain kinds of inclusions may serve to protect cells and to assist in the clearance of these toxic species [8-17]. There is strong evidence that the progressive spread of pathology in patients could be due to the cell-to-cell propagation of protein misfolding and aggregation [18, 19]. However, the formation of protein inclusions is also indicative of dysregulated protein homeostasis (proteostasis) [20, 21] and the inability of the cell to properly monitor, refold or degrade non-native proteins.

Given the genetic and functional heterogeneity of ALS, the development of high content assays that can measure multiple phenotypic features in cellular ALS models and extract rich, descriptive information of responses to candidate therapeutic compounds and modifiers of ALS gene toxicity will be valuable. To address this need, we have developed an experimental system with HCA capacity that can be used to extract multiplexed phenotypic data from cellular ALS models. In the present work we have tested this system to examine proteostasis capacity in cellular models of SOD1- and CCNF-linked ALS.

## 2. MATERIALS AND METHODS

### 2.1. Plasmids

All plasmids are detailed in the Supplementary Material.

### 2.2. Cell culture

NSC-34 cells [22] were maintained in 10% (v/v) fetal bovine serum (FBS; Bovogen Biologicals) in Dulbecco’s Modified Eagle’s Medium/Ham’s Nutrient Mixture F-12 (DMEM/F-12). Cells were seeded into either 8-well µ-Slides (Ibidi) for confocal microscopy or 96-well plates (Greiner Bio-One) for imaging using the Cellomics ArrayScan VTI imaging platform (Thermo Scientific). After overnight incubation at 37 °C under 5% CO_2_/95% air, cells were either single-transfected or triple-transfected (detailed in Tables S1 and S2, supplementary information) using Lipofectamine 2000 (Invitrogen) according to manufacturer’s instructions. For triple transfections, plasmids were used at a 1: 1: 1 ratio. All transfection conditions were carried out in quadruplicate. For experiments involving proteasome inhibition, MG132 was solubilised in DMSO at 20 mM and subsequently diluted to 5 µM in 10% (v/v) FBS in DMEM/F-12. The prepared solution was added to cells 30 h post-transfection and incubated for 18 h.

### 2.3. Confocal microscopy

Localisation of each fusion protein in transfected NSC-34 cells was characterised by imaging using a TCS SP5 II confocal microscope with a 63× oil-immersion objective lens (Leica Microsystems). Imaging was carried out 48 h post-transfection.

### 2.4. High-content analysis

NSC-34 cells were imaged either live or fixed (in 4% (w/v) paraformaldehyde for 20 min at room temperature) using the 20× objective of a Cellomics ArrayScan VTI imaging platform (Thermo Scientific). All assays described are appropriate for either live or fixed cell imaging. Fluorescence of ECFP-, EGFP- and tdT/mCherry-fusion proteins was imaged using excitation filters of 386 nm, 485 nm and 549 nm, respectively. Phase contrast and fluorescent images from 20 fields of view per well were acquired, with image analysis parameters optimised using the SpotDetector V4 BioApplication in HCS Studio (Thermo Scientific) summarised in Figure 1 (further detail in Figure S1). Primary object identification was based on expression of H2B-ECFP in channel 1, while GFP or tdTomato/mCherry-fusion proteins were detected in channels 2 and 3.

**Figure 1.**
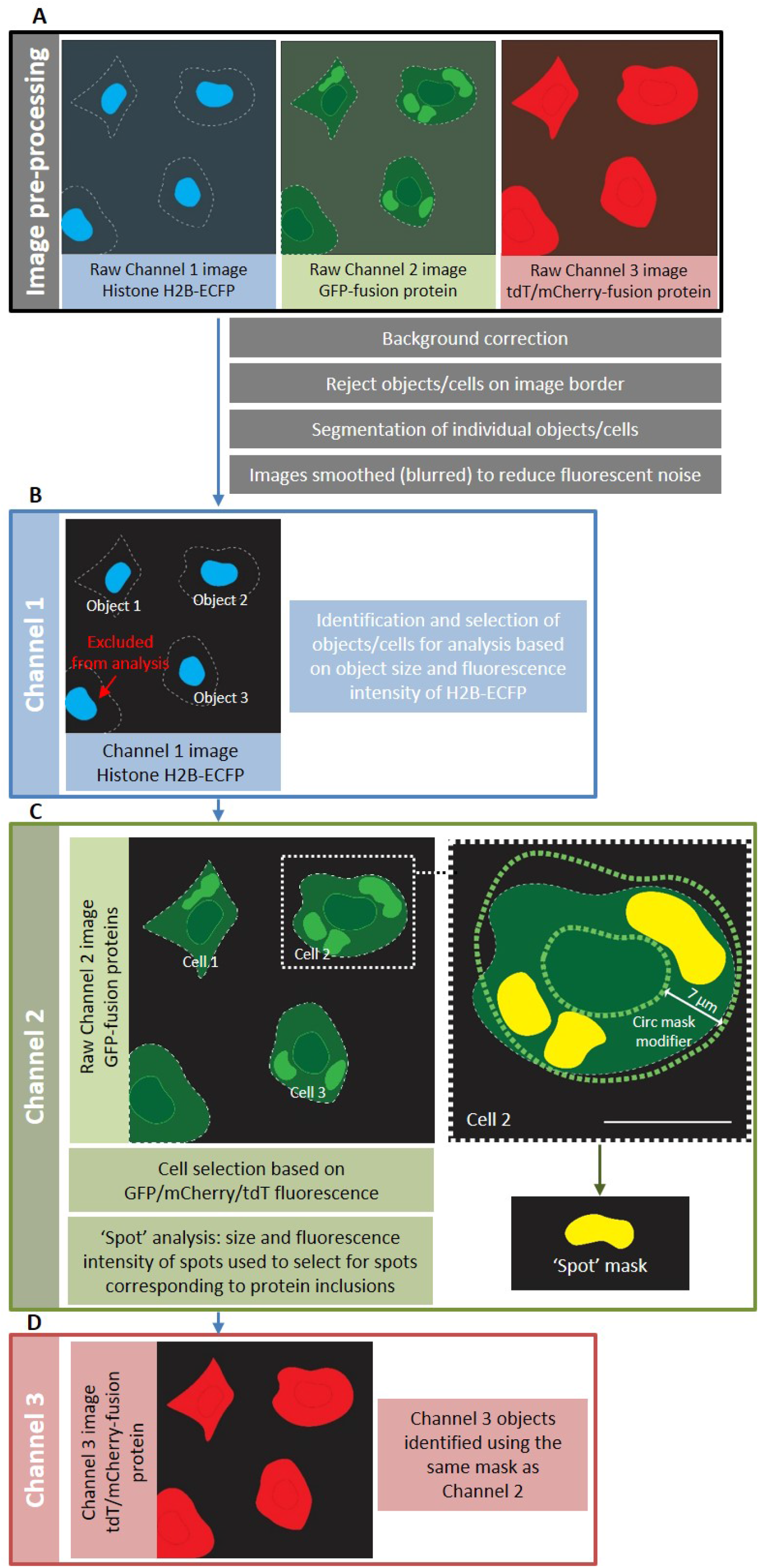
Schematic of High Content Analysis (HCA) image processing and analysis optimisation. To analyse the fluorescence intensity of EGFP-/tGFP- and tdTomato (tdT)/mCherry-fusion proteins and quantify protein inclusions containing EGFP-/tGFP-fusion proteins in NSC-34 cells, we deployed the Spot Detector BioApplication designed to analyse fluorescent foci in cells. Optimisation was carried out using images of NSC-34 cells triple-transfected to express H2B-ECFP, either SOD1^WT^-EGFP, SOD1^A4V^-EGFP, TDP-43^WT^-tGFP TDP-43^M337V^-tGFP, FUS^WT^-tGFP, FUS^R495X^-tGFP, FUS^R521G^-tGFP or EGFP alone and mCherry alone. Cells were imaged at 48 h post-transfection using a 20× objective lens. (A) Raw images from Channels 1 (H2B-ECFP), 2 (EGFP-/tGFP-fusion proteins) and 3 (tdTomato/mCherry-fusion proteins) were first pre-processed to remove background fluorescence, exclude cells positioned on the border of each image from analysis and distinguish individual cells (‘object’ segmentation). Channel 1 images were additionally smoothed (blurred) to help reduce fluorescent noise that could lead to the false inclusion of image artefacts in subsequent analyses. (B) Biological ‘objects’, in this case cells, were identified using nuclear-localised H2B-ECFP fluorescence in Channel 1 images. To select viable transfected cells for analysis and exclude image artefacts, dead cells and cell debris, cells were selected based on the size and fluorescence intensity of their ECFP-fluorescent nuclei. (C) The relevant measures for GFP fluorescence intensity and fluorescent foci were measured in Channel 2 within a circular analysis mask that expanded the mask derived in Channel 1. The green circular mask indicates cells selected for analysis, while yellow masks indicate fluorescent foci/’spots’ selected for analysis. To detect and analyse fluorescent foci corresponding to protein inclusions, upper and lower limits for size and fluorescence intensity were set. (D) Channel 3 objects were identified using the same mask as Channel 2.

## 3. RESULTS

### 3.1. Characterisation of cellular ALS models

Given the extraordinary molecular heterogeneity of ALS, we developed a suite of ALS models representing genetically diverse fALS aetiologies in NSC-34 cells expressing EGFP-/tGFP-/mCherry-fusions of mutant SOD1, TDP-43, FUS, CCNF, UBQLN2, OPTN, VCP or VAPB. The genetic mutations for these models were selected after careful consideration of the mutations that segregate with ALS; *SOD1*^*A4V*^ [23], *TARDBP*^*M337V*^ [24], *FUS*^*R495X*^, *FUS*^*R521G*^ [3, 4], *CCNF*^*S621G*^ [25], *UBQLN2*^*P497H*^ [26], *OPTN*^*E478G*^ [27], *VAPB*^*P56S*^ [28], *VCP*^*R159H*^ and *VCP*^*R191Q*^ [29]. Prior to using these NSC-34 models in an HCA format, we first characterised the localisation, toxicity, and solubility of each WT and mutant fusion protein (Figure S3-S7). Importantly, EGFP and mCherry were diffusely distributed throughout the cell and did not aggregate (Figure S2, a, i and b, i). It was also confirmed that the expression of EGFP or mCherry-alone had no effect on cell viability (Figure S2, a, ii and b, ii).

### 3.2. Toxicity of mutant SOD1, TDP-43, FUS and CCNF

Live cell imaging of cells expressing SOD1^WT^-EGFP and SOD1^A4V^-EGFP to monitor cell population growth showed that the numbers of cells expressing SOD1^WT^-EGFP and EGFP-alone steadily increased, while the numbers of cells expressing SOD1^A4V^-EGFP increased at a slower rate (Figure 2, a, iii). At 48 h post-replating there were significantly lower numbers of cells expressing SOD1^A4V^-EGFP than SOD1^WT^-EGFP (*p* = 0.0151) (Figure 2, a, iv). This indicates that the overexpression of SOD1^A4V^-EGFP caused toxicity.

**Figure 2.**
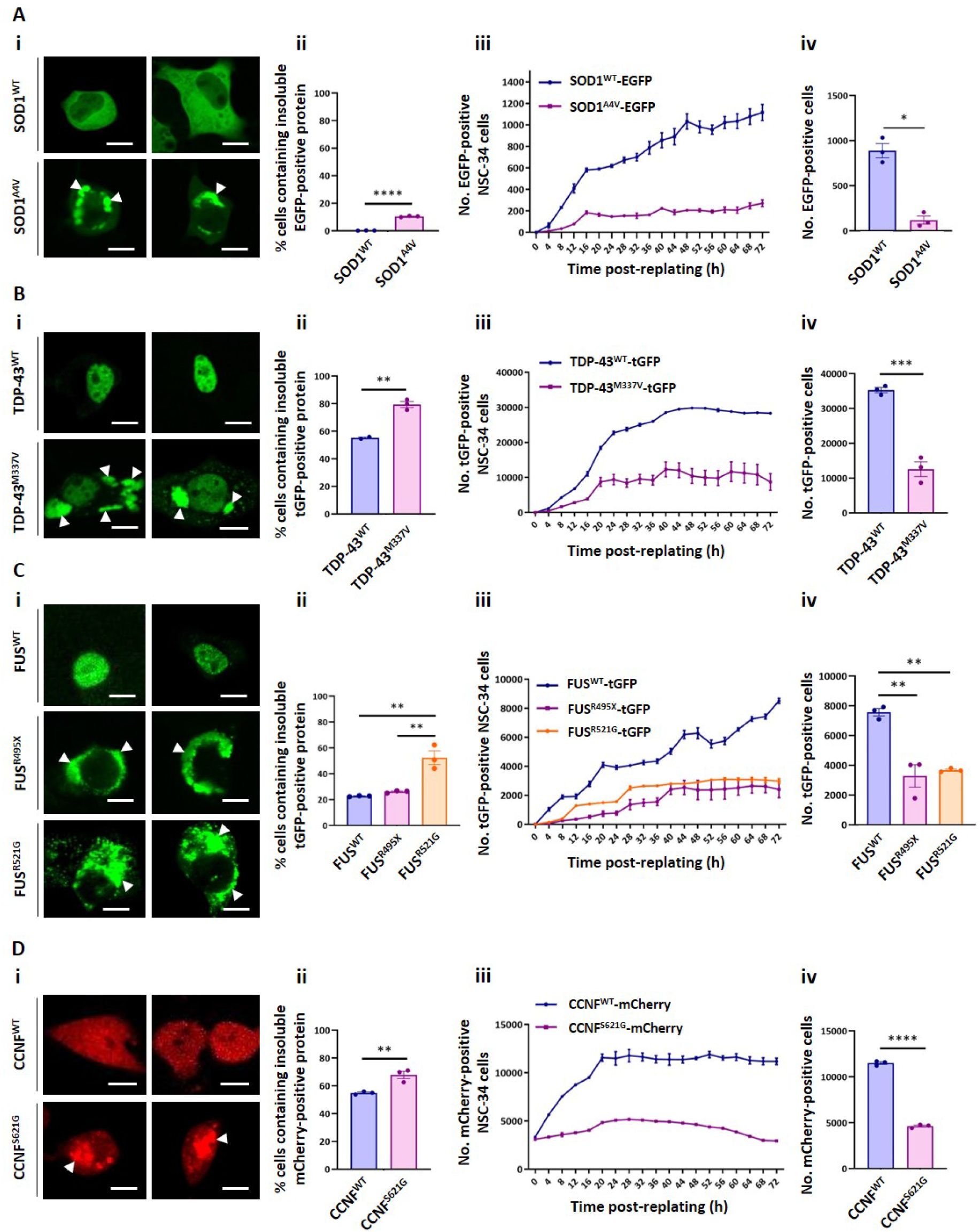
Characterising the localisation patterns and intracellular solubility of ALS-associated SOD1^A4V^, TDP-43^M337V^, FUS^R495X^, FUS^R521G^ and CCNF^S621G^. NSC-34 cells were transiently transfected with (A) SOD1^WT^-EGFP or SOD1^A4V^-EGFP, (B) TDP-43^WT^-tGFP or TDP-43^M337V^-tGFP, (C) FUS^WT^-tGFP, FUS^R495X^-tGFP or FUS^R521G^-tGFP or (D) CCNF^WT^-mCherry or CCNF^S621G^-mCherry. After 48 h, transfected cells were either (i) imaged using a Leica TCS SP5 II confocal microscope, (ii) imaged on an IncuCyte® ZOOM, followed by incubation with 0.03% (w/v) saponin in PBS for 10 min at room temperature, before being imaged again on the IncuCyte or (iii and iv) imaged in an IncuCyte® ZOOM over 72 h. (i) Representative images from confocal microscopy, with white arrow heads indicating inclusions formed by the EGFP/tGFP/mCherry-fusion proteins. Scale bars represent 10 µm. (ii) Cells were transfected in quadruplicate, and the data presented is the mean ± SEM of the percentage of transfected NSC-34 cells containing insoluble EGFP-/tGFP-/mCherry-positive protein following permeabilisation with saponin. (iii) Numbers of EGFP/tGFP/mCherry-positive transfected cells over 72 h and (iv) the mean ± SEM numbers of transfected cells at 48 h post-replating, in triplicate wells of cells. Differences between the means were determined using Student’s t test or one-Way ANOVA followed by Tukey’s Multiple Comparison Test. * indicates *p* < 0.05, ** indicates *p* < 0.01, *** indicates *p* < 0.001 and **** indicates *p* < 0.0001.

A similar trend was observed for cells expressing TDP-43, FUS or CCNF. Comparison of the mean numbers of GFP/RFP-positive transfected cells at 72 h post-transfection showed that there was a significantly greater number of cells expressing WT proteins than cells expressing mutants (TDP-43^M337V^-tGFP, *p* < 0.001; FUS^R495X^-tGFP, *p* = 0.0014; FUS^R521G^-tGFP, *p* = 0.0023; CCNF^S621G^-mCherry, *p* < 0.0001) (Figure 2, b-d, iv).

### 3.3. Localisation and aggregation of mutant SOD1, TDP-43, FUS and CCNF

Inclusions of ALS-associated proteins are generally 2-20 µm in diameter in both human post-mortem tissue [27, 30-33] and in cell culture models [34-39]. A size minimum of 2 µm was thus established as suitable for categorising fluorescent foci as inclusions. The foci formed by SOD1^A4V^-EGFP, TDP-43^M337V^-tGFP, FUS^R495X^-tGFP, FUS^R521G^-tGFP and CCNF^S621G^-mCherry were manually examined in images of cells and were consistently measured to be larger than 2 µm (Figure 2, a, i-d, i).

While SOD1^WT^-EGFP was observed to have a relatively even distribution throughout the cytoplasm and within nuclei, in a proportion of cells SOD1^A4V^-EGFP formed multiple large inclusions in the cytoplasm (Figure 2, a, i). Saponin-permeabilisation showed that there was no fluorescent signal from cells overexpressing SOD1^WT^-EGFP following permeabilisation, indicating that SOD1^WT^-EGFP remained soluble in all cells that were imaged (Figure 2, a, ii). In accordance with the confocal microscopy data, a significantly greater percentage of cells (10.38 ± 0.27%) expressing SOD1^A4V^-EGFP remained EGFP-positive following permeabilisation (*p* < 0.0001), indicating that SOD1^A4V^-EGFP was present in an insoluble, non-diffusable form in a proportion of cells.

Imaging transfected cells using confocal microscopy, it was observed that TDP-43^WT^ and FUS^WT^ remained localised to cell nuclei, while the TDP-43 and FUS mutants mislocalised to the cytoplasm and formed large aggregates and smaller foci, as is observed in ALS patient tissue (Figure 2, b, i and c, i) [3, 4, 24]. Although TDP-43^WT^-tGFP was not observed to mislocalise and accumulate into cytoplasmic inclusions when cells were examined using confocal microscopy, TDP-43^WT^-tGFP was found to remain inside 55.22 ± 0.69% of transfected cells after plasma membrane permeabilisation (Figure 2, b, ii). However, a significantly greater percentage of cells expressing TDP-43^M337V^-tGFP were tGFP-positive following permeabilisation (79.41 ± 2.18%) compared to cells expressing TDP-43^WT^-tGFP (*p* = 0.0035). TDP-43^WT^-tGFP that was bound to immobile elements or granules may not have been released when cells were incubated with saponin solution. In this case, the saponin-permeabilisation assay may not be appropriate for assaying the formation of TDP-43 cytoplasmic inclusions. Similarly, while FUS^WT^-tGFP was not observed to mislocalise and accumulate into cytoplasmic inclusions when transfected cells were examined using confocal microscopy, the saponin-permeabilisation assay quantified that FUS^WT^-tGFP remained in 22.72 ± 0.25% of cells following incubation with saponin solution (Figure 2, c, ii). Moreover, the percentage of cells expressing FUS^R495X^-tGFP that remained tGFP-positive following permeabilisation (26.15 ± 0.56%) was similar to that of cells expressing FUS^WT^-tGFP. However, there was a significantly greater percentage of cells expressing FUS^R521G^-tGFP that remained tGFP-positive (52.4 ± 5.29%) compared to both cells expressing FUS^WT^-tGFP and cells expressing FUS^R495X^-tGFP (FUS^WT^-tGFP, *p* = 0.0012; FUS^R495X^-tGFP, *p* = 0.0023). As noted above, when cells expressing the FUS-tGFP constructs were examined using confocal microscopy, there was extensive formation of small foci (< 2 µm) and large aggregates by both FUS mutants (Figure 2, c, i). Thus, the similar percentages of cells expressing FUS^WT^-tGFP and FUS^R495X^-tGFP that remained tGFP-positive after saponin-permeabilisation compared to the marked differences in their localisation patterns indicates that the saponin-permeabilisation assay may not be appropriate for measuring the formation of insoluble cytoplasmic mutant FUS-tGFP inclusions.

Since the original identification of the *CCNF*^*S621G*^ mutation in ALS and frontotemporal dementia patients [25], the localisation patterns of CCNF^S621G^ in motor neurons have not been investigated in detail. However, Lee *et al*. [40] observed CCNF^S621G^ localised to inclusion-like structures while CCNF^WT^ displayed diffuse distribution. We observed mCherry-tagged CCNF^WT^ fluorescence with diffuse distribution in all imaged cells (Figure 2, d, i). In contrast, CCNF^S621G^-mCherry formed into large amorphous aggregates ranging from 5 to > 10 µm. Interestingly, saponin-permeabilisation revealed that both CCNF^WT^-mCherry and CCNF^S621G^-mCherry formed extensively into insoluble structures, with > 50% of both cells transfected with CCNF^WT^-mCherry cells and CCNF^S621G^-mCherry cells containing insoluble mCherry-positive protein (Figure 2, d, ii). Nevertheless, there were significantly more CCNF^S621G^-mCherry-expressing cells containing insoluble mCherry-positive protein (67.84 ± 2.61%) than there were of CCNF^WT^-mCherry cells (54.89 ± 0.64%) (*p* = 0.0085).

### 3.4. HCA assay for proteostasis stress in cells expressing SOD1^A4V^ and CCNF^S621G^

The optimised HCA SpotDetector BioApplication was used to compare the effect of SOD1 and CCNF variant overexpression on the ability of the cellular protein quality control network to prevent aggregation of conformationally-destabilised, aggregation-prone mutants of firefly luciferase (Fluc) [41-45]. It was reasoned that reductions in protein quality control network capacity would lead to increased aggregation of the EGFP-tagged Fluc mutants [41].

We detected > 90% of ECFP- and EGFP-positive cells (green circular masks). ‘Spot’ masks (yellow masks) were observed only on large foci with high fluorescence intensity, corresponding to the inclusion size and fluorescence intensity cut-offs established during assay optimisation.

Proteasome inhibition of cells expressing mCherry alone confirmed that increased proteome stress results in increased aggregation of the Fluc-EGFP isoforms. Cells expressing mCherry that were treated with MG132 developed significantly greater numbers of Fluc^WT^-EGFP (*p* < 0.0001), Fluc^SM^-EGFP (*p* < 0.0001) and Fluc^DM^-EGFP (*p* < 0.0001) aggregates compared to untreated cells (Figure 4, a, i). MG132 treatment resulted in significantly higher numbers of Fluc^WT^-EGFP aggregates than Fluc^DM^-EGFP aggregates (*p* = 0.0153). Fluc^WT^-EGFP aggregates were significantly smaller (*p* = 0.002) and more brightly fluorescent (*p* < 0.0001) than Fluc^DM^-EGFP aggregates (Figure 4, b, i and c, i). Aggregates of the Fluc-EGFP isoforms were also detected in cells expressing SOD1-tdT, with a significant increase in the numbers of aggregates formed in cells expressing SOD1^A4V^-tdT compared to SOD1^WT^-tdT (Fluc^WT^-EGFP, *p* = 0.0331; Fluc^SM^-EGFP, *p* = 0.0061; Fluc^DM^-EGFP, *p* = 0.0042) (Figure 4, a, ii). There were also increases in the mean size of Fluc^DM^-EGFP aggregates (*p* = 0.0430) and fluorescence intensity of aggregates of Fluc^WT^-EGFP (*p* < 0.0001), Fluc^SM^-EGFP (*p* < 0.0001) and Fluc^DM^-EGFP (*p* < 0.0001) in cells expressing SOD1^A4V^-tdT compared to SOD1^WT^-tdT (Figure 4, b, ii and c, ii). Interestingly, aggregation of Fluc^SM^-EGFP and Fluc^DM^-EGFP was as extensive in cells expressing CCNF^WT^-mCherry as those expressing CCNF^S621G^-mCherry (Figure 4, a, iii). Whilst aggregates of Fluc^WT^-EGFP were also detected, there were significantly lower numbers of cells containing them compared to the numbers of cells containing aggregates of Fluc^SM^-EGFP (*p* < 0.001) and Fluc^DM^-EGFP (*p* < 0.0001), both in cells expressing CCNF^WT^-mCherry and cells expressing CCNF^S621G^-mCherry. There was also the same trend in the size of aggregates of the Fluc-EGFP isoforms in cells expressing CCNF^WT^-mCherry and those expressing CCNF^S621G^-mCherry, with significantly larger aggregates of Fluc^DM^-EGFP formed than aggregates of Fluc^WT^-EGFP (CCNF^WT^-mCherry, *p* = 0.0395; CCNF^S621G^-mCherry, *p* = 0.0328) (Figure 4, b, iii). There was no difference in the mean fluorescence intensity of the Fluc-EGFP aggregates of the Fluc-EGFP isoforms between cells expressing CCNF^WT^-mCherry and cells expressing CCNF^S621G^-mCherry (Figure 4, c, iii).

**Figure 3.**
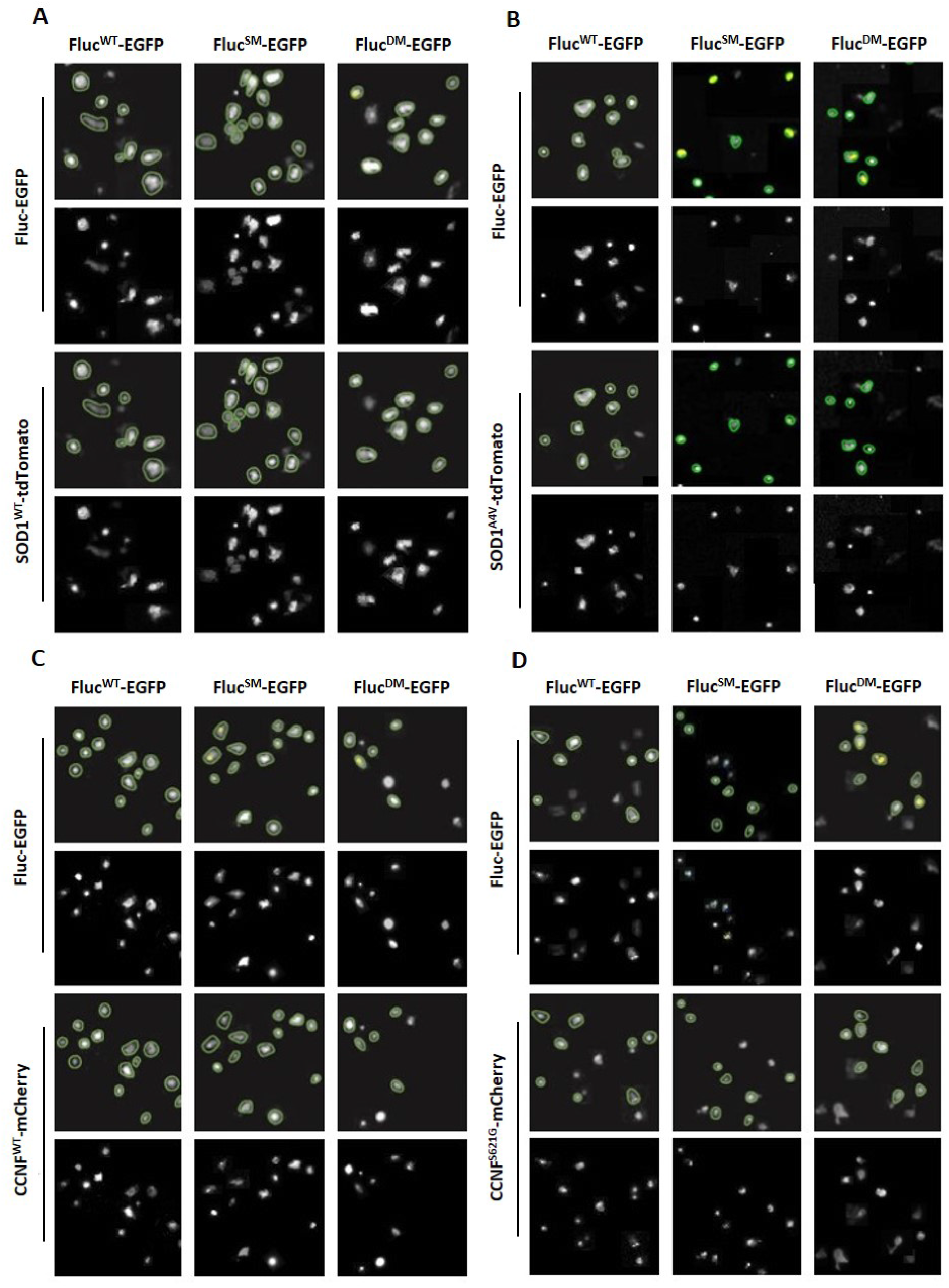
Optimised HCA SpotDetector BioApplication identifies and analyses transfected cells and Fluc-EGFP foci. Representative Cellomics® ArrayScan™ VTI images showing masks (first and third rows of each panel) used to identify and select NSC-34 cells co-transfected with either (A) SOD1^WT^-tdTomato, (B) SOD1^A4V^-tdTomato, (C) CCNF^WT^-mCherry or (D) CCNF^S621G^-mCherry and Fluc^WT^-EGFP, Fluc^SM^-EGFP or Fluc^DM^-EGFP. Cells were imaged at 48 h post-transfection. Green circular masks indicate cells selected for analysis, yellow masks indicate ‘spots’ selected for analysis, representing aggregates. Images were acquired using a 20× objective lens.

**Figure 4.**
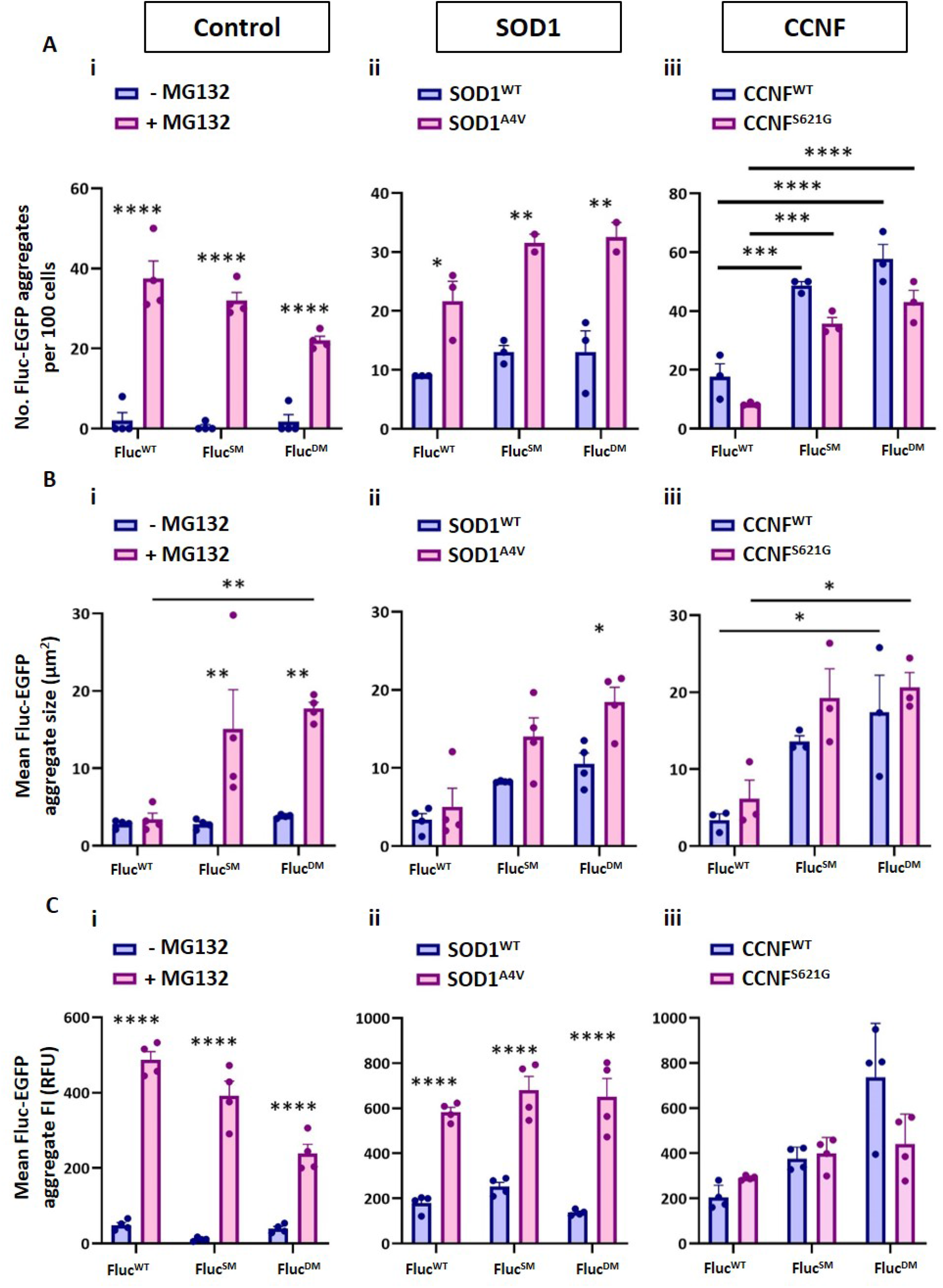
Firefly luciferase mutants report on chaperone network activity in NSC-34 cells expressing SOD1 and CCNF. (A) Numbers of Fluc-EGFP aggregates per 100 transfected cells, (B) mean size of Fluc-EGFP aggregates (µm^2^) and (C) mean fluorescence intensity (FI) of Fluc-EGFP aggregates imaged at 48 h post-transfection in NSC-34 cells expressing (i) mCherry alone ± treatment with 5 µM MG132, (ii) SOD1^WT^-tdTomato or SOD1^A4V^-tdTomato or (iii) CCNF^WT^-mCherry or CCNF^S621G^-mCherry. Treatment with MG132 was carried out at 30 h post-transfection. For mock treatment, 5 µM DMSO was instead added to cells. Graphs represent the mean ± SEM from quadruplicate wells of cells in n = 1 experiment. Differences between the means were determined using one-Way ANOVA followed by Tukey’s Multiple Comparison Test. * indicates *p* < 0.05, ** indicates *p* < 0.01, *** indicates *p* < 0.001, **** indicates *p* < 0.0001.

## 4. DISCUSSION

The genetic heterogeneity of ALS distinguishes it from most other neurodegenerative diseases, which can be linked to a limited number of pathogenic mechanisms and phenotypes. The ALS research field would benefit greatly from the use of experimental systems with HCA capacity to help navigate through the complexity of ALS. To address this, we have developed an HCA methodology to use with cellular ALS models. The overall objective of this work was to develop a system that could be used to collect descriptive phenotypic data from cellular ALS models that would enable (1) characterisation of the inclusion formation pathways of different ALS-associated proteins, and (2) the use of diverse markers of proteome stress and motor neuron dysfunction to assess potential therapeutic compounds and genetic modifiers of ALS disease mechanisms and toxicity.

The NSC-34 models of ALS generated here were examined for the localisation, mobility and solubility of the fusion proteins. The aim of these studies was to establish disease phenotypes that could be used in an experimental system with HCA capacity for further studies into disease mechanisms, and potentially for evaluation of candidate therapeutics. This HCA experimental system was generated through optimising the SpotDetector BioApplication to investigate reductions in cellular protein folding/re-folding capacity caused by WT and mutant SOD1 and CCNF. Analysis was facilitated by co-expression of conformationally-destablised Fluc-EGFP mutants [41] with WT and mutant SOD1 and CCNF. We hypothesised that dysregulation of proteostasis mechanisms that may be exacerbated by ALS-associated mutations would overload cellular proteostasis capacity, resulting in inability of the cellular pool of molecular chaperones to prevent aggregation of the Fluc-EGFP mutants. The optimised SpotDetector BioApplication enabled quantification of the numbers, mean size and fluorescence intensity of aggregates formed by the Fluc-EGFP isoforms.

The ability of the Fluc-EGFP mutants to report on proteome stress was confirmed through proteasome inhibition of cells expressing mCherry alone. In MG132-treated cells, Fluc^WT^-EGFP aggregates that formed were smaller than the aggregates formed by Fluc^SM^-EGFP and Fluc^DM^-EGFP, indicating that less of the WT protein misfolded and accumulated into aggregates. Without exogenous proteome stress induced by proteasome inhibition, there was negligible aggregation of the Fluc-EGFP isoforms, demonstrating that they were able to report on increased proteome stress.

The data from the present work demonstrates that the optimised Fluc-EGFP HCA assay is able to report on reduced activity of the chaperone network resulting from the expression of SOD1^A4V^ as previously reported [46-48]. The overexpression of CCNF^WT^-mCherry caused the same extent of Fluc-EGFP aggregation as CCNF^S621G^-mCherry, indicating that mutant CCNF did not differentially affect chaperone activity compared to CCNF^WT^. CCNF is an important protein in the ubiquitin-proteasome system, as a mediator of protein ubiquitylation [49]. Ubiquitylation of target proteins is altered in cells expressing mutant CCNF, causing aberrant accumulation of ubiquitylated proteins and consequent stress on the proteostasis network [25]. The data obtained from the Fluc-EGFP HCA assay developed in the present work suggests that proteostasis disruption caused by mutant CCNF does not involve impairment of the protein folding/re-folding activity of chaperones.

The Fluc-EGFP isoforms were designed to act as sensors of cellular protein folding/re-folding capacity that would themselves have minimal biological impact in most of the commonly used cellular and animal models [41]. In the present work it was demonstrated that they are suitable for use in an HCA assay format to report on disruptions in the activity of the cellular chaperone network. In future work it would be useful to optimise an HCA assay that utilises changes in luminescence activity of the Fluc-EGFP isoforms [41] as an additional measure.

In addition to the use of this Fluc-EGFP HCA assay to examine cellular models of SOD1^A4V^ and CCNF^S621G^, it would be useful in future work to utilise this assay to examine the cellular models of mutant TDP-43, FUS, UBQLN2, OPTN, VAPB and VCP generated in this work. Beyond establishing ALS-associated mutant proteins that impair the activity of the chaperone network in cells, this HCA assay could have potential for application in studies to screen for drugs that ameliorate chaperone activity impairment.

## Funding & Acknowledgements

J.J.Y. was supported by a University of Wollongong Professorship in Neurodegenerative Diseases, and by an National Health and Medical Research Council, Australia Dementia Teams Grant (1095215) and Investigator Grant (1194872).

## CRediT author statement

**Isabella A. Lambert-Smith**: Conceptualisation, Methodology, Validation, Formal analysis, Investigation, Data curation, Writing – Original Draft, Visualisation. **Justin J. Yerbury:** Conceptualisation, Methodology, Formal analysis, Investigation, Resources, Writing – Review & Editing, Visualisation, Supervision, Project administration, Funding acquisition. **Darren N. Saunders:** Conceptualisation, Methodology, Formal analysis, Investigation, Resources, Writing – Review & Editing, Visualisation, Supervision, Project administration, Funding acquisition.

