## Supplemental Material for "High-content analysis of proteostasis capacity in cellular models of amyotrophic lateral sclerosis (ALS)"

**Supplementary Material**

Isabella A. Lambert-Smith[1,2], Justin J. Yerbury[1,2], Darren N. Saunders[1]

1. Illawarra Health and Medical Research Institute, University of Wollongong, Wollongong, NSW 2522 Australia

2. School of Chemistry and Molecular Bioscience, University of Wollongong, Wollongong, NSW 2522 Australia

#### Supplementary Materials and Methods

##### Plasmids

The pEGFP-N1 expression vector containing human wild-type (WT) *SOD1* (SOD1$^{WT}$-EGFP) and *SOD1* altered by the disease-associated Ala4Val (A4V) mutation (SOD1$^{A4V}$-EGFP) were constructed as previously described [1]. pEGFP-C3 containing human OPTN$^{WT}$ (OPTN$^{WT}$-EGFP), OPTN$^{Q398X}$ (OPTN$^{Q398X}$-EGFP) and OPTN$^{E478G}$ (OPTN$^{E478G}$-EGFP) were provided by Assoc. Prof. Julie Atkin (Macquarie University, NSW, Australia). Mammalian expression constructs comprised of the pCMV6-AC-tGFP vector containing human WT *TARDBP* (TDP-43$^{WT}$-tGFP), *FUS/TLS (*FUS$^{WT}$-tGFP), *VAPB* (VAPB$^{WT}$-tGFP), *UBQLN2* (UBQLN2$^{WT}$-tGFP) and *VCP* (VCP$^{WT}$-tGFP) were obtained from OriGene Technologies (MD, U.S.A.). To obtain constructs containing ALS-associated mutations in each gene, sequences were designed in-house according to mutant sequences described in the literature, and site-directed mutagenesis was outsourced to GenScript. The following mutants were generated: TDP-43$^{M337V}$-tGFP; FUS$^{R495X}$-tGFP; FUS$^{R521G}$-tGFP; VAPB$^{P56S}$-tGFP; UBQLN2$^{P497H}$-tGFP; VCP$^{R159H}$-tGFP; VCP$^{R191Q}$-tGFP. pcDNA3.1(+) containing human SOD1$^{WT}$ or SOD1$^{A4V}$, each with C-terminal tdTomato fluorescent tags (SOD1$^{WT}$-tdT and SOD1$^{A4V}$-tdT), were custom-cloned using GenScript. Plasmid constructs containing human WT *CCNF* (Cyclin F; CCNF$^{WT}$-mCherry) and CCNF$^{S621G}$ (CCNF$^{S621G}$-mCherry) in pmCherry-C1 were donated by Prof. Ian Blair (Macquarie University, NSW, Australia). The pBOS-H2B-ECFP-N1 construct, generated as described by Leung *et al* [2], was generously provided by Prof. Angus Lamond (University of Dundee, UK).

The pCIneo mammalian expression construct containing sequences encoding C-terminally EGFP-tagged WT firefly luciferase (Fluc$^{WT}$-EGFP) and two conformationally destabilised mutants (containing a single mutation (SM), Fluc$^{SM}$-EGFP or two mutations (double mutant; DM), Fluc$^{DM}$-EGFP) were generously provided by Prof. Franz-Ulrich Hartl (Max Planck Institute of Biochemistry, Planegg, Germany). For detailed information on the design and construction of the destabilised Fluc-EGFP mutant plasmids see Gupta *et al.* [3].

##### Cell culture, transient transfections and treatment with proteasome inhibitor

For confocal microscopy experiments, NSC-34 cells were transfected with $SOD1^{WT}$-EGFP, $SOD1^{A4V}$-EGFP, TDP-$43^{WT}$-tGFP, TDP-$43^{M337V}$-tGFP, $FUS^{WT}$-tGFP, $FUS^{R495X}$-tGFP, $FUS^{R521G}$-tGFP, $CCNF^{WT}$-mCherry, $CCNF^{S621G}$-mCherry, the pEGFP-N1 vector (EGFP alone) or the pmCherry-C1 vector (mCherry alone). For optimisation of the parameters used for automated imaging and image analysis using the Thermo Scientific™ Cellomics® ArrayScan™ VTI High Content Screening microscope, NSC-34 cells were triple-transfected as detailed in Table S1.

**Table S1. Plasmid combinations used in triple transfections in preparation for HCA microscopy optimisation.** Triple-transfected NSC-34 cells were used for preliminary experiments to optimise the parameters used for automated imaging and image analysis using a Cellomics® ArrayScan™ VTI HCS microscope.

| Plasmid 1 | Plasmid 2 | Plasmid 3 |
|---|---|---|
| H2B-ECFP | $SOD1^{WT}$-EGFP | mCherry alone |
| | $SOD1^{A4V}$-EGFP | |
| | TDP-$43^{WT}$-tGFP | |
| | TDP-$43^{M337V}$-tGFP | |
| | $FUS^{WT}$-tGFP | |
| | $FUS^{R495X}$-tGFP | |
| | $FUS^{R521G}$-tGFP | |
| | $VAPB^{WT}$-tGFP | |
| | $VAPB^{P56S}$-tGFP | |
| | $OPTN^{WT}$-EGFP | |
| | $OPTN^{E478G}$-EGFP | |
| | $UBQLN2^{WT}$-tGFP | |
| | $UBQLN2^{P497H}$-tGFP | |
| | $VCP^{WT}$-tGFP | |
| | $VCP^{R159H}$-tGFP | |
| | $VCP^{R191Q}$-tGFP | |

**Table S2. Plasmid combinations used in triple transfections of NSC-34 cells in preparation for Fluc-EGFP experiments.**

| Plasmid 1 | Plasmid 2 | Plasmid 3 | Additional treatment |
|---|---|---|---|
| H2B-ECFP | $SOD1^{WT}$-tdT | $Fluc^{WT}$-EGFP | No treatment |
| | | $Fluc^{SM}$-EGFP | |

| | | | |
|---|---|---|---|
| | | Fluc$^{DM}$-EGFP | |
| | | EGFP alone | |
| H2B-ECFP | SOD1$^{A4V}$-tdT | Fluc$^{WT}$-EGFP | No treatment |
| | | Fluc$^{SM}$-EGFP | |
| | | Fluc$^{DM}$-EGFP | |
| | | EGFP alone | |
| H2B-ECFP | CCNF$^{WT}$-mCherry | Fluc$^{WT}$-EGFP | No treatment |
| | | Fluc$^{SM}$-EGFP | |
| | | Fluc$^{DM}$-EGFP | |
| | | EGFP alone | |
| H2B-ECFP | CCNF$^{S621G}$-mCherry | Fluc$^{WT}$-EGFP | No treatment |
| | | Fluc$^{SM}$-EGFP | |
| | | Fluc$^{DM}$-EGFP | |
| | | EGFP alone | |
| H2B-ECFP | mCherry alone | Fluc$^{WT}$-EGFP | ± MG132 treatment |
| | | Fluc$^{SM}$-EGFP | |
| | | Fluc$^{DM}$-EGFP | |
| | | EGFP alone | |

**Confocal image analysis and inclusion characterisation**

To characterise the localisation patterns of each EGFP-/tGFP- and mCherry-tagged ALS-associated protein in transfected NSC-34 cells and ensure that they corresponded with the patterns recorded in the literature, the images acquired by confocal microscopy were manually examined. To inform this characterisation, previous peer-reviewed studies revealing images of post-mortem spinal cord tissue from ALS patients were used to establish the morphology and size range of inclusions positive for mutant SOD1, TDP-43, FUS, VAPB, OPTN, UBQLN2 and VCP. This enabled a size minimum of 2 µm to be established for categorising fluorescent foci as inclusions as opposed to associations of the fluorescent ALS proteins with other cellular structures, granules (e.g stress or transport) or other irrelevant fluorescent debris and artefacts.

**IncuCyte® ZOOM live cell imaging and analysis**

IncuCyte® ZOOM live cell imaging and analysis was used to carry out two different measures in transfected NSC-34 cells. One of the experiments involved monitoring the relative population growth of transfected cells (EGFP-positive cells) over 72 h. The net cell population growth (rate of cell division – rate of cell death) was used as a measure of cell viability, on the basis that toxicity caused by the overexpression of the mutant ALS gene would impair cellular ability to proliferate and/or result in the loss of cells relative to control cells overexpressing the WT gene or EGFP/mCherry alone. NSC-34 cells were transfected in 6-well plates and were re-plated at 24 h post-transfection at uniform cell density ($5 \times 10^4$ cells/mL) in 200 µL/well DMEM/F-12/FBS into 96-well plates and imaged using the 10× objective lens of an IncuCyte® ZOOM over 72 h. To quantify the numbers of viable transfected cells, a processing definition in the IncuCyte® ZOOM software was optimised to identify and count transfected cells based on a minimum level of EGFP/mCherry fluorescence, and viable cells based on a minimum size. Non-transfected controls were used to account for and exclude background fluorescence and fluorescent artefacts, cell debris and dying cells.

The second experiment quantified cells containing inclusions using a modification of a saponin-permeabilisation protocol described by Pokrishevsky *et al.* (2018) [4]. Saponin is a mild, cholesterol-chelating detergent that creates pores in the plasma membrane of cells [5]. These pores allow soluble intracellular proteins to diffuse out of the cell, while trapping insoluble proteinaceous structures. NSC-34 cells were transfected with each expression construct in quadruplicate in 96-well plates, and after 48 h were imaged using an IncuCyte® ZOOM. These pre-permeabilisation images were used to measure the numbers of cells that were transfected (EGFP-/mCherry-positive) in each well. After imaging, cells were incubated with 0.03% (w/v) saponin in PBS for 10 min at RT to create pores in the cells' plasma membranes. Saponin-treated cells were then imaged again to measure the numbers of cells in which non-diffusable, insoluble EGFP-/mCherry-positive material remained following permeabilisation. The percentage of transfected cells containing insoluble EGFP-/mCherry-fusion proteins were then calculated using the following formula:

$$\%\ transfected\ cells\ containing\ insoluble\ fusion\ proteins = \left(\frac{Number\ of\ GFP\ or\ mCherry\ positive\ cells\ post\ permeabilisation}{Number\ of\ GFP\ or\ mCherry\ positive\ cells\ pre\ permeabilisation}\right) \times 100$$

Equation S1.

For all IncuCyte® experiments, the mean ± SEM was calculated across quadruplicate replicates, and used for statistical analyses.

**High-content analysis**

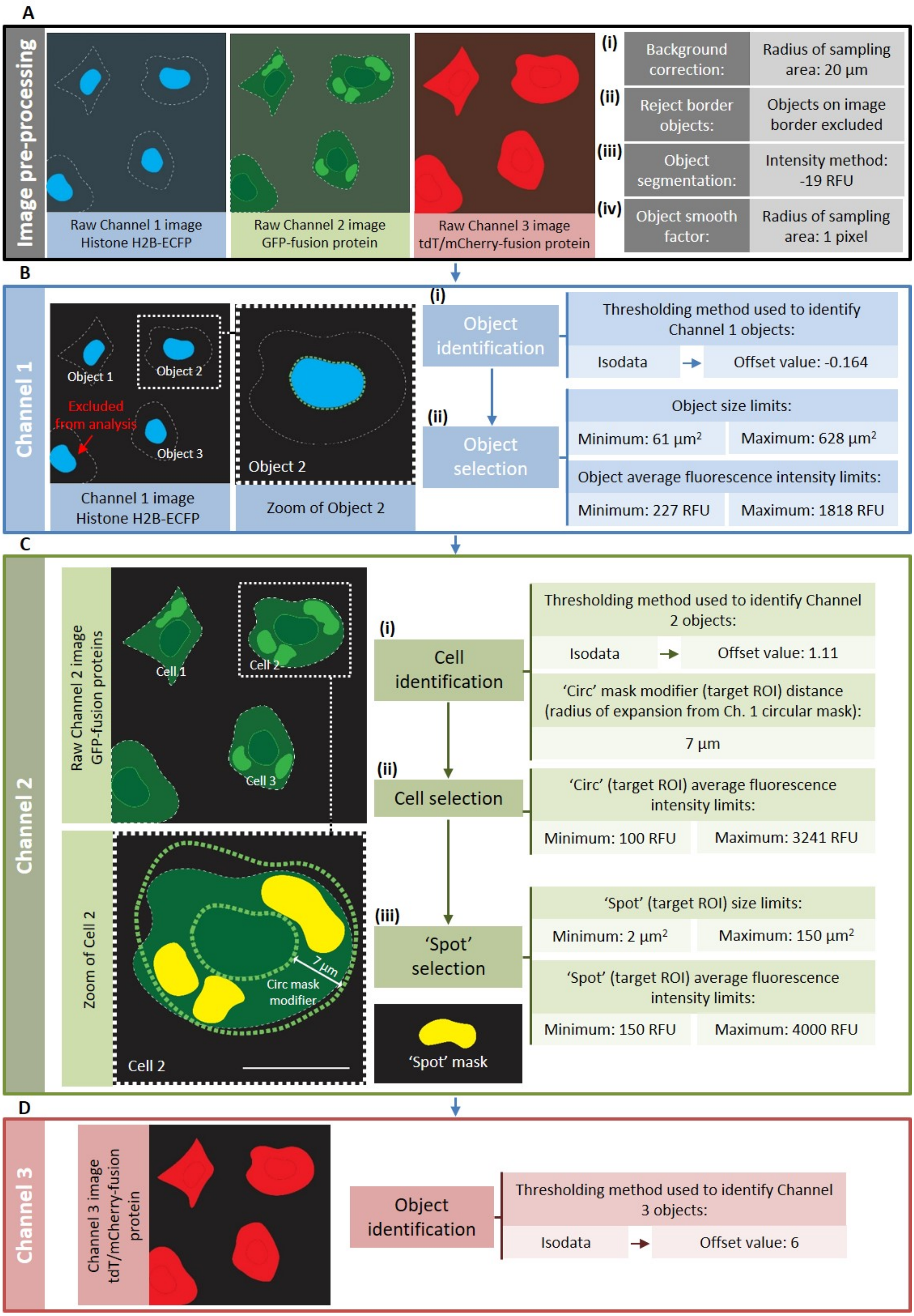


**Figure S1. Schematic of Cellomics® ArrayScan™ VTI High Content Screening (HCS) image processing and analysis optimisation using Thermo Scientific™ HCS Studio™ software.** To analyse the fluorescence intensity of EGFP-/tGFP- and tdTomato (tdT)/mCherry-fusion proteins and quantify protein inclusions containing EGFP-/tGFP-fusion proteins in NSC-34 cells, an image analysis algorithm designed to analyse fluorescent foci in cells, termed the Cellomics® Spot Detector BioApplication, was optimised using the Thermo Scientific™ HCS Studio™ software. Optimisation was carried out using images of NSC-34 cells triple-transfected to express H2B-ECFP, either $SOD1^{WT}$-EGFP, $SOD1^{A4V}$-EGFP, TDP-$43^{WT}$-tGFP TDP-$43^{M337V}$-tGFP, $FUS^{WT}$-tGFP, $FUS^{R495X}$-tGFP, $FUS^{R521G}$-tGFP or EGFP alone and (3) mCherry alone. Cells were imaged at 48 h post-transfection using the 20× objective lens of a Cellomics® ArrayScan™ VTI HCS microscope. (A) Raw images from Channels 1 (H2B-ECFP), 2 (EGFP-/tGFP-fusion proteins) and 3 (tdTomato/mCherry-fusion proteins) were first pre-processed to (i) remove background fluorescence using a low-pass filtration method (local background around each pixel is calculated, with the radius of the area sampled adjusted as determined by the user), (ii) exclude cells positioned on the border of each image from analysis and (iii) distinguish individual cells; 'object' segmentation. An object segmentation method based on fluorescence intensity was used, which separates objects/cells that are touching based on fluorescence intensity peaks of each pixel. Cells expressing H2B-ECFP in their nucleus exhibit a single, high intensity peak localised in the nucleus. Setting this parameter, ObjectSegmentationCh1, to a negative value sets the segmentation method to the Intensity method. The absolute value selected governs the minimum relative height of the intensity peak to be used for segmentation. A value of -19 relative fluorescence units (RFU) was found to optimally separate touching fluorescent cells. (iv) Channel 1 images were additionally smoothed (blurred) to help reduce fluorescent noise that could lead to the false inclusion of image artefacts in subsequent analyses. (B) Biological 'objects', in this case cells, were identified using nuclear-localised H2B-ECFP fluorescence in Channel 1 images. (i) For detection of ECFP-fluorescent nuclei, a fluorescence intensity threshold was set using the Isodata method, which derives the threshold from the distribution of pixel intensities in each image. (ii) To select viable transfected cells for analysis and exclude image artefacts, dead cells and cell debris, cells were selected based on the size and fluorescence intensity of their ECFP-fluorescent nuclei. (C) The relevant measures for GFP fluorescence intensity and fluorescent foci were measured in Channel 2 (i) within a circular analysis mask that expanded the mask derived in Channel 1 by 7 µm. The green circular mask indicates cells selected for analysis, while yellow masks indicate fluorescent foci/'spots' selected for analysis. A fluorescence intensity threshold was set for Channel 2 using the Isodata method. An offset value of 1.11 enabled the image processing algorithm to correctly identify cells expressing the GFP-fusion proteins. (ii) To ensure that cells with particularly low expression of GFP-fusion genes were not included for analysis, GFP fluorescence intensity limits for CircAvgIntenCh2 were set to a minimum intensity of 100 RFU and a maximum intensity of 3241 RFU. (iii) To detect and analyse fluorescent foci corresponding to protein inclusions, as established using confocal microscopy, a spot area minimum of 2 $\mu m^2$ and maximum of 150 $\mu m^2$ was set. As inclusions are accumulations of high concentrations of proteins, the concentrated fluorescence of GFP-tagged proteins in inclusions allows them to be detected based on high fluorescence intensity. A spot average fluorescence intensity minimum of 150 RFU was found to detect appropriate foci corresponding to GFP-positive inclusions. (D) Channel 3 objects were identified using the same mask as Channel 2, after establishing the fluorescence intensity threshold using the Isodata method with an offset value of 6.

### RESULTS

#### Characterisation of cellular ALS models

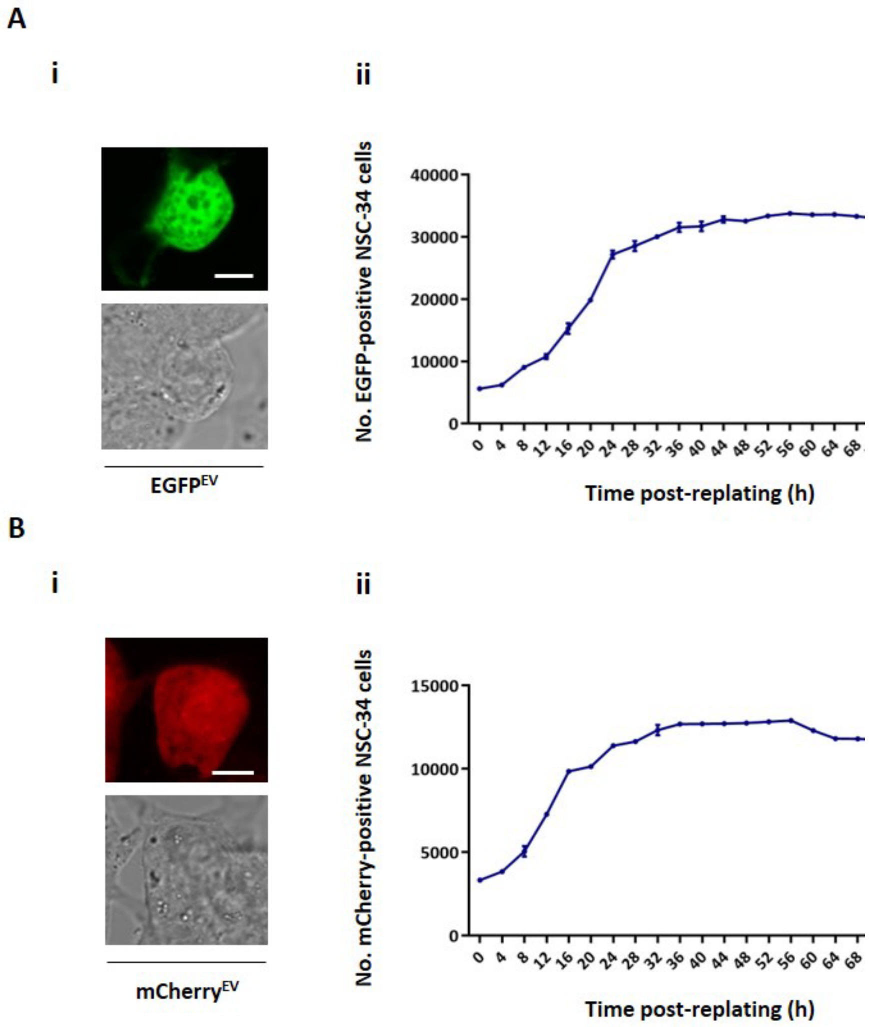


**Figure S2. Localisation patterns of EGFP and mCherry alone, and cell population growth over time of NSC-34 cells expressing EGFP or mCherry alone.** Live cell imaging of NSC-34 cells transiently transfected with (A) the pEGFP-N1 vector (EGFP alone) or (B) the pmCherry-C1 vector (mCherry alone). (A, i and B, i) Representative images of NSC-34 cells transiently transfected with (A) the pEGFP-N1 vector (EGFP alone) or (B) the pmCherry-C1 vector (mCherry alone). Cells were imaged using a Leica TCS SP5 II confocal microscope at 48 h post-transfection. Scale bars represent 10 µm. (A, ii and B, ii) To measure viability of NSC-34 cells transiently transfected to express EGFP or mCherry alone, cells were imaged in an IncuCyte® ZOOM and numbers of transfected cells (EGFP- or mCherry-positive cells) were monitored over 68 h. Graphs represent the mean ± SEM of the numbers of EGFP- or mCherry-positive transfected cells over 68 h.

**Toxicity of mutant UBQLN2, OPTN, VAPB and VCP**

Assaying cell population growth using live cell imaging over 72 h, expression of UBQLN2$^{P497H}$-tGFP and of UBQLN2$^{P525S}$-tGFP was found to cause reductions in cell population growth rates compared to the expression of UBQLN2$^{WT}$-tGFP (UBQLN2$^{P497H}$-tGFP, $p < 0.0001$; UBQLN2$^{P525S}$-tGFP, $p < 0.001$) (Figure S3, b). Interestingly, expression of the UBQLN2$^{P497H}$-tGFP mutant was observed to cause greater inhibition of cell population growth compared to the UBQLN2$^{P525S}$-tGFP mutant, with UBQLN2$^{P497H}$-tGFP cells exhibiting slower population growth rates ($p = 0.0275$) and dropping down to significantly lower numbers by 72 h post-transfection ($p = 0.0264$) than UBQLN2$^{P525S}$-tGFP cells (Figure S3, c, i).

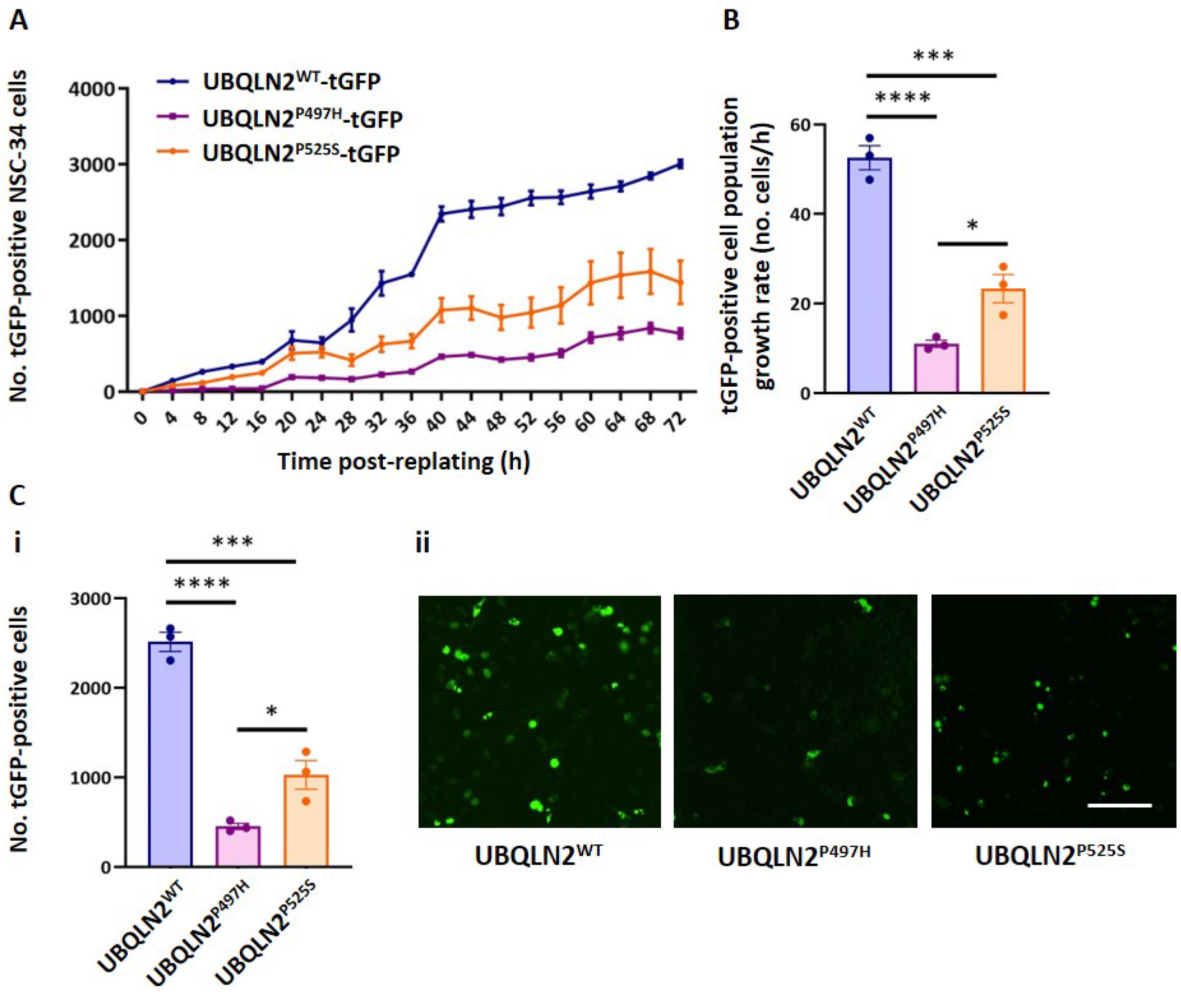


**Figure S3. ALS-associated UBQLN2$^{P497H}$ and UBQLN2$^{P525S}$ cause toxicity in NSC-34 cells.** NSC-34 cells were transiently transfected with UBQLN2$^{WT}$-tGFP, UBQLN2$^{P497H}$-tGFP or UBQLN2$^{P525S}$-tGFP and imaged in an IncuCyte® ZOOM over 72 h. Graphs represent the mean ± SEM of (A) numbers of tGFP-positive transfected cells over 72 h, (B) population growth rates of transfected cells and (C, i) numbers of transfected cells at 48 h post-replating, in triplicate wells of cells. (C, ii) Representative IncuCyte images of NSC-34 cells at 48 h post-replating, from which the graph in C, i, was derived. Scale bar represents 150 µm. Differences between the means were determined using one-Way ANOVA followed by Tukey's Multiple Comparison Test. * indicates $p < 0.05$, *** indicates $p < 0.001$, **** indicates $p < 0.0001$.

In NSC-34 cells expressing the VAPB-tGFP constructs, a clear reduction was observed in the population growth of cells expressing VAPB$^{P56S}$-tGFP compared to cells expressing VAPB$^{WT}$-tGFP (Figure S4). Although numbers of transfected cells were low due to low transfection efficiency, total cell density was high, increasing from ~70% to ~95% over the 72 h imaging period (data not shown). There was thus no potential effect of low total cell density on the proliferation or loss of the VAPB-tGFP-expressing cells. Comparison of the numbers of VAPB-tGFP cells in the first 20 h of imaging confirmed that VAPB$^{P56S}$-tGFP significantly reduced the population growth rate of cells ($p$ = 0.0203) (Figure S4, b). The numbers of tGFP-positive cells expressing VAPB$^{P56S}$-tGFP at 48 h post-replating were significantly lower than cells expressing VAPB$^{WT}$-EGFP ($p$ < 0.0001) (Figure S4, c, i), further highlighting the toxicity caused by expression of the P56S mutation.

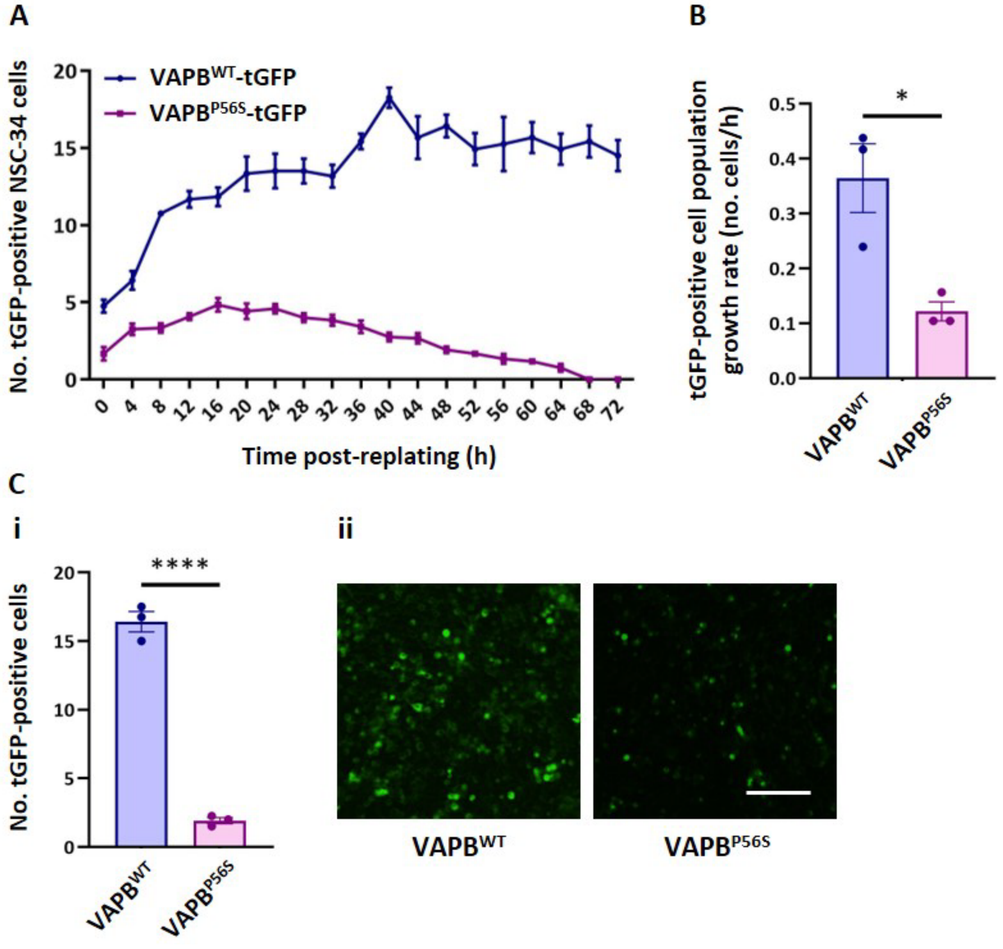


**Figure S4. ALS-associated VAPB$^{P56S}$ causes toxicity in NSC-34 cells.** NSC-34 cells were transiently transfected with VAPB$^{WT}$-tGFP or VAPB$^{P56S}$-tGFP and imaged in an IncuCyte® ZOOM over 72 h. Graphs represent the mean ± SEM of (A) numbers of tGFP-positive transfected cells over 72 h, (B) population growth rates of transfected cells and (C, i) numbers of transfected cells at 48 h post-replating, in triplicate wells of cells. (C, ii) Representative IncuCyte images of NSC-34 cells at 48 h post-replating, from which the graph in C, i, was derived. Scale bar represents 150 μm. Differences between the means were determined using Student's t tests. * indicates $p < 0.05$, **** indicates $p < 0.0001$.

The expression of OPTN$^{E478G}$-EGFP reduced the population growth rate of transfected cells compared to cells expressing OPTN$^{WT}$-EGFP (Figure S5). In the first 20 h of imaging, the population growth rate of OPTN$^{E478G}$-EGFP-expressing cells was significantly slower than that of OPTN$^{WT}$-EGFP expressing cells ($p$ = 0.0139) (Figure S5, b). Comparison of the numbers of transfected cells at 48 h post-replating showed that there were significantly more viable OPTN$^{WT}$-EGFP-expressing cells than of OPTN$^{E478G}$-EGFP-expressing cells ($p$ < 0.001) (Figure S5, c, i), demonstrating that overexpression of the E478G mutant caused considerable toxicity in NSC-34 cells.

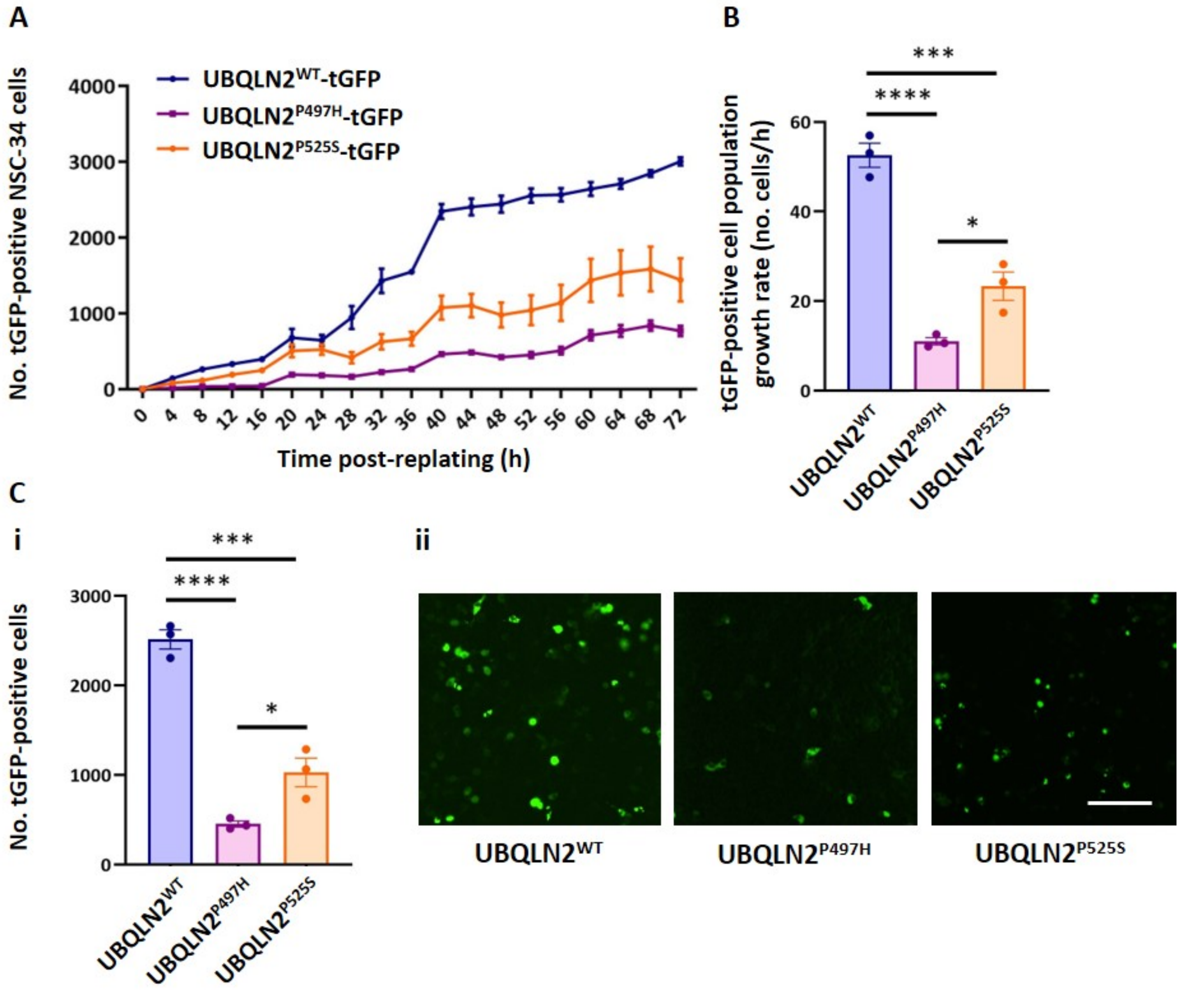


**Figure S5. ALS-associated OPTN$^{E478G}$ causes toxicity in NSC-34 cells.** NSC-34 cells were transiently transfected with OPTN$^{WT}$-EGFP or OPTN$^{E478G}$-EGFP and imaged in an IncuCyte® ZOOM over 72 h. Graphs represent the mean ± SEM of (A) numbers of EGFP-positive transfected cells over 72 h, (B) population growth rates of transfected cells and (C, i) numbers of transfected cells at 48 h post-replating, in triplicate wells of cells. (C, ii) Representative IncuCyte images of NSC-34 cells at 48 h post-replating, from which the graph in C, i, was derived. Scale bar represents 150 µm. Differences between the means were determined using Student's t tests. * indicates $p < 0.05$, *** indicates $p < 0.001$.

Assaying the numbers of NSC-34 cells expressing the VCP-tGFP constructs over 72 h, it was observed that both mutants significantly reduced the population growth rate of transfected cells relative to $VCP^{WT}$-tGFP (Figure S6). The population growth rate of cells expressing $VCP^{R191Q}$-tGFP was notably slower than that of cells expressing $VCP^{R159H}$-tGFP ($p = 0.004$), and 48 h after replating cells the numbers of viable $VCP^{R191Q}$-tGFP-expressing cells were much lower than the numbers of $VCP^{R159H}$-tGFP cells ($p = 0.0012$) (Figure S6, b and c, i).

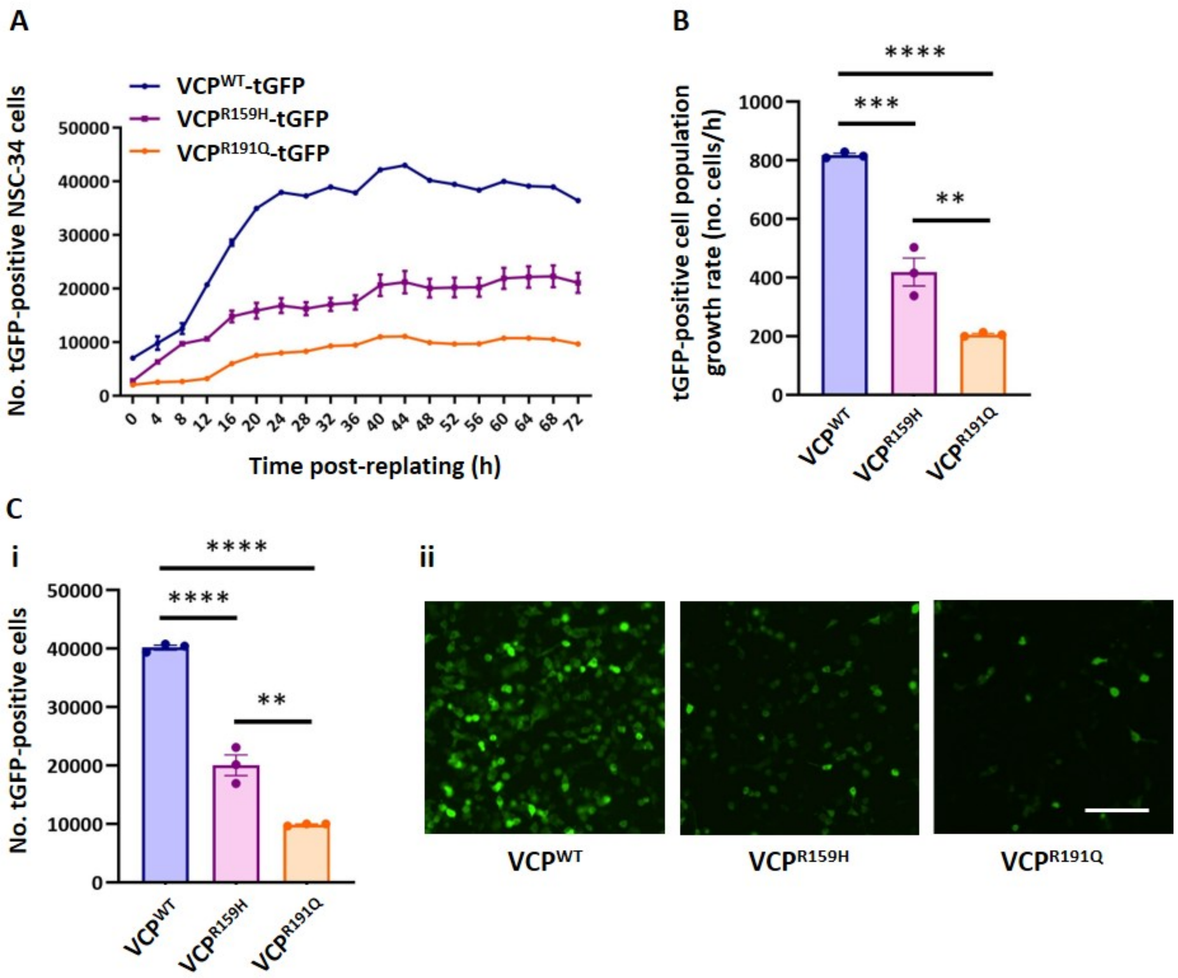


**Figure S6. ALS-associated $VCP^{R159H}$ and $VCP^{R191Q}$ cause toxicity in NSC-34 cells.** NSC-34 cells were transiently transfected with $VCP^{WT}$-tGFP, $VCP^{R159H}$-tGFP or $VCP^{R191Q}$-tGFP and imaged in an IncuCyte® ZOOM over 72 h. Graphs represent the mean ± SEM of (A) numbers of tGFP-positive transfected cells over 72 h, (B) population growth rates of transfected cells and (C, i) numbers of transfected cells at 48 h post-replating, in triplicate wells of cells. (C, ii) Representative IncuCyte images of NSC-34 cells at 48 h post-replating, from which the graph in C, i, was derived. Scale bar represents 150 µm. Differences between the means were determined using one-Way ANOVA followed by Tukey's Multiple Comparison Test. ** indicates $p < 0.01$, *** indicates $p < 0.001$ and **** indicates $p < 0.0001$.

**Characterisation of mutant UBQLN2, OPTN, VAPB and VCP solubility, localisation and aggregation**

The saponin-permeabilisation assay was used to examine the mobility and solubility of UBQLN2$^{P497H}$-tGFP, UBQLN2$^{P525S}$-tGFP, OPTN$^{E478G}$-EGFP, VAPB$^{P56S}$-tGFP, VCP$^{R159H}$-tGFP and VCP$^{R191Q}$-tGFP in NSC-34 cells. In cells expressing the UBQLN2-tGFP constructs it was found that UBQLN2$^{WT}$-tGFP had a degree of immobility in cells, with 37% of transfected cells quantified to contain tGFP-positive protein following permeabilisation of their plasma membranes (Figure S7, a). However, the two UBQLN2 mutants, UBQLN2$^{P497H}$-tGFP and UBQLN2$^{P525S}$-tGFP, remained inside significantly more cells following permeabilisation; 65.1% ($p$ = 0.002) and 68.1% ($p$ = 0.0012), respectively. Interestingly, the saponin-permeabilisation assay quantified that ~35% of both cells expressing OPTN$^{WT}$-EGFP and cells expressing OPTN$^{E478G}$-EGFP contained insoluble EGFP-positive protein following permeabilisation (Figure S7, b). As OPTN$^{WT}$ can associate with proteins of the Golgi apparatus [6, 7], a proportion of OPTN$^{WT}$-EGFP may have been bound to the Golgi and thus immobile and unable to diffuse out of cells following saponin treatment. Further experimentation is needed to determine the nature of this post-permeabilisation cell-bound OPTN$^{WT}$-EGFP. Through saponin permeabilisation of cells transfected with the VCP-tGFP constructs it was quantified that ~9.8% of cells transfected with VCP$^{WT}$-tGFP retained the tGFP-tagged protein following membrane permeabilisation (Figure S7, c). However, both VCP$^{R191Q}$-tGFP and VCP$^{R159H}$-tGFP remained cell-associated in significantly greater percentages of transfected cells (VCP$^{R191Q}$-tGFP, $p$ = 0.0222; VCP$^{R159H}$-tGFP, $p$ = 0.0111), indicating that both mutants were relatively less mobile and had reduced solubility compared to VCP$^{WT}$-tGFP.

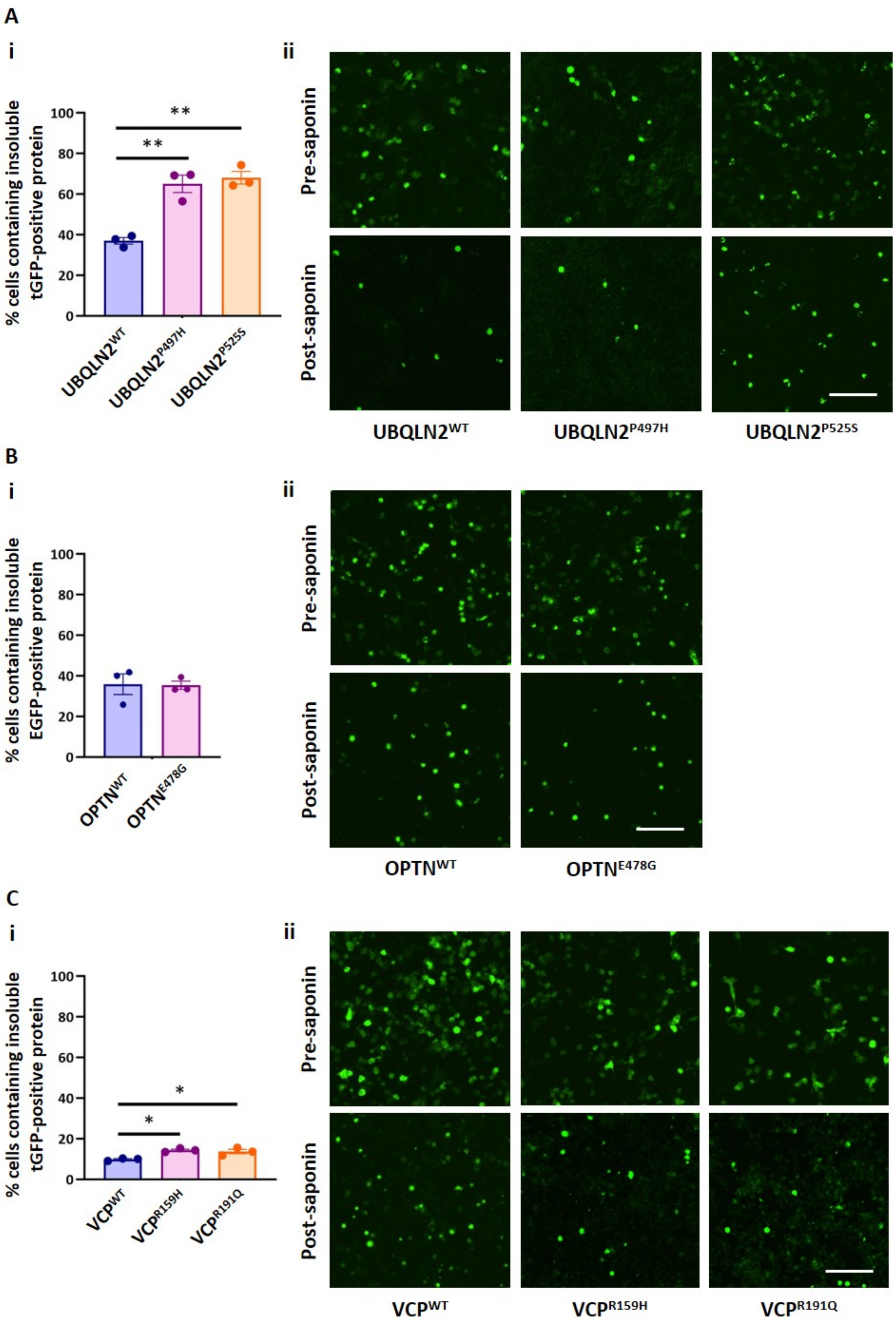
A
i
ii
% cells containing insoluble tGFP-positive protein
100
80
60
40
20
0
**
**
UBQLN2WT
UBQLN2P497H
UBQLN2P525S
Pre-saponin
Post-saponin
B
i
ii
% cells containing insoluble EGFP-positive protein
OPTNWT
OPTNE478G
C
i
ii
% cells containing insoluble tGFP-positive protein
*
*
VCPWT
VCPR159H
VCPR191Q

**Figure S7. Analysis of the release of EGFP-/tGFP-fusion mutant UBQLN2, OPTN and VCP from NSC-34 cells following saponin-permeabilisation.** NSC-34 cells were transiently transfected with (A) UBQLN2WT-tGFP, UBQLN2P497H-tGFP or UBQLN2P525S-tGFP, (B) OPTNWT-EGFP or OPTNE478G-EGFP or (C) VCPWT-tGFP, VCPR159H-tGFP or VCPR191Q-tGFP. After 48 h, cells were imaged on an IncuCyte® ZOOM, followed by incubation with 0.03% (w/v) saponin in PBS for 10 min at room temperature, before being imaged again on the IncuCyte. (A, i, B, i and C, i) Cells were transfected in quadruplicate, and the data presented in each graph is the mean ± SEM of the percentage of transfected NSC-34 cells containing insoluble EGFP-/tGFP-positive protein following permeabilisation with saponin. Differences between the means were determined using either Student's t tests or one-Way ANOVA followed by Tukey's Multiple Comparison Test. * indicates $p < 0.05$, ** indicates $p < 0.01$. (A, ii, B, ii and C, ii) Representative IncuCyte images of NSC-34 cells prior to permeabilisation with saponin (pre-saponin) and immediately following permeabilisation with saponin (post-saponin), from which the graphs in A, i, B, i and C, i) were derived. Scale bars represent 150 µm.

Unfortunately, transfections with the VAPB-tGFP constructs for the IncuCyte-based assays resulted in considerably low transfection efficiencies, as can be seen in the low numbers of tGFP-positive transfected cells in Figure S4. The numbers of tGFP-positive cells at 48 h post-transfection detected pre- and post-permeabilisation with saponin for VAPB$^{P56S}$-tGFP were thus too low for analysis. This saponin-treatment experiment also may not have been appropriate for VAPB, as it is an integral membrane protein and thus in its native localisation in cells is immobile and unable to freely diffuse out of permeabilised cells.
